# Sex change in aquarium systems establishes the New Zealand spotty wrasse (*Notolabrus celidotus*) as a temperate model species for the investigation of sequential hermaphroditism

**DOI:** 10.1101/2020.08.28.271973

**Authors:** A Goikoetxea, S Muncaster, EV Todd, PM Lokman, HA Robertson, CE De Farias e Moraes, EL Damsteegt, NJ Gemmell

**Author notes:** These authors contributed equally to this article (A Goikoetxea, S Muncaster). Current affiliation: ^1^MARBEC Univ Montpellier, CNRS, Ifremer, IRD, Palavas-Les-Flots, France. Corresponding Author: Simon Muncaster, School of Science, University of Waikato Private Bag 3105, Hamilton 3240, New Zealand.

## Abstract

The stunning sexual transformation commonly triggered by age, size or social context in some fishes is one of the best examples of phenotypic plasticity thus far described. To date our understanding of this process is dominated by studies on a handful of subtropical and tropical teleosts, often in wild settings because sex change has been challenging to achieve in captivity. Here we have established the protogynous New Zealand spotty wrasse, *Notolabrus celidotus*, as a temperate model for the experimental investigation of sex change. Captive fish were induced to change sex using either aromatase inhibition or manipulation of social groups. Complete transition from female to male occurred over 60 days and time-series sampling was used to quantify changes in hormone production, gene expression and gonadal cellular anatomy using radioimmunoassay, nanoString nCounter mRNA and histological analyses, respectively. Early-stage decreases in plasma 17β-estradiol (E2) concentrations or gonadal aromatase (*cyp19a1a*) expression were not detected in spotty wrasse, despite these being commonly associated with the onset of sex change in subtropical and tropical protogynous (female-to-male) hermaphrodites. In contrast, expression of the masculinising factor *amh* (anti-Müllerian hormone) increased during early sex change, implying a potential role as a proximate trigger for masculinisation. Expression of male-related genes responsible for androgen production *cyp11c1* and *hsd11b2* increased from mid sex change. Gonadal expression of the glucocorticoid and mineralocorticoid receptors *nr3c1* and *nr3c2*, putative mediators of the stress hormone cortisol, increased in late stages of sex change. Collectively, these data provide a foundation for the spotty wrasse as a temperate teleost model to study sex change and cell fate in vertebrates.

**Summary statement:** The spotty wrasse, *Notolabrus celidotus*, is a new temperate model for the study of vertebrate sex change, this work characterises endocrine and genetic markers based on laboratory induced sex change.

## Introduction

For most vertebrates, sexual fate is genetically determined and remains fixed throughout life. However, for many teleost fishes sex is more plastic (Avise and Mank, 2009). The complete sex change that sequentially hermaphroditic fishes undergo during their reproductive lives, while unique among vertebrates, is taxonomically widespread across the teleost tree (Avise and Mank, 2009; Godwin, 2009; Munday et al., 2006). The direction and process of sex change differ greatly among teleosts (Gemmell et al., 2019; Goikoetxea et al., 2017; Todd et al., 2016) and appear to have evolved multiple times (Avise and Mank, 2009). The cellular and molecular processes that underpin sex change are slowly being determined for a few focal species (Todd et al., 2019). However, the extent to which these processes might be conserved and reused across the teleosts to achieve sex change is poorly understood (Ortega-Recalde et al., 2020).

Changes in the social environment and community structure often cue the timing of sex change (Reavis and Grober, 1999; Solomon-Lane et al., 2013; Sprenger et al., 2012). Species where manipulation of social structure can readily induce natural sex change are convenient models to understand the mechanistic drivers of this transformation. More broadly, these species present *in vivo* opportunities to examine cell fate pathways, neuro-endocrine plasticity, genetic and epigenetic regulation of life-history trajectory and reproductive status.

Current sex change research mostly focuses on tropical and subtropical models within the Labridae (Godwin et al., 1996; Kojima et al., 2008; Lamm et al., 2015; Liu et al., 2017; Nakamura et al., 1989; Nozu et al., 2009; Ohta et al., 2003), Serranidae (Alam et al., 2008; Bhandari et al., 2003; Bhandari et al., 2005; Chen et al., 2020b; Li et al., 2007) and Gobiidae (Kroon and Liley, 2000; Kroon et al., 2005; Maxfield and Cole, 2019). Unfortunately, such manipulations have frequently been limited to the wild because sex change in laboratory settings has proven challenging for many of these species. In addition, few studies have focused on temperate fishes that experience strong reproductive seasonality and a protracted period of sex change. These species, arguably, offer an extended window of graded change in which to tease out fine-scale modulation of physiological and molecular drivers.

The New Zealand (NZ) spotty wrasse, *Notolabrus celidotus*, is an endemic protogynous (female-to-male sex change) temperate zone (35° S – 47° S) labrid that is well suited to laboratory studies. These small (< 26 cm) fish are abundant and easily caught on shallow reefs and in harbours around the NZ coastline. They have dimorphic initial phase (IP) and terminal phase (TP) colour morphs, characteristic of most wrasses (Choat, 1965; Jones, 1980). As in other wrasses, two male sexual strategies exist with IP sneaker males displaying female mimicry and behaviourally dominant TP males establishing defended breeding territories (Fig. 1). Reproduction in the spotty wrasse peaks in the austral spring but the exact timing varies with latitude (Jones, 1980), the NZ coastline spanning around 12 ° of latitude. This physically hardy species has a wide thermal tolerance (approximately 8 – 25° C) and adapts well to captivity and tolerates experimental manipulation. Sexually mature fish will spawn in captivity and sex change is induced in IP fish through the manipulation of social structure (Thomas et al., 2019). This proclivity to complete natural sex change under laboratory conditions is of particular significance as other model species such as the bluehead wrasse (*Thalassoma bifasciatum*) adapt poorly to captivity leading to most sex change experiments being done in wild populations (Liu et al., 2015; Thomas et al., 2019; Todd et al., 2019). Collectively, these attributes make spotty wrasse an excellent biological model to study sex change.

**Figure 1.**
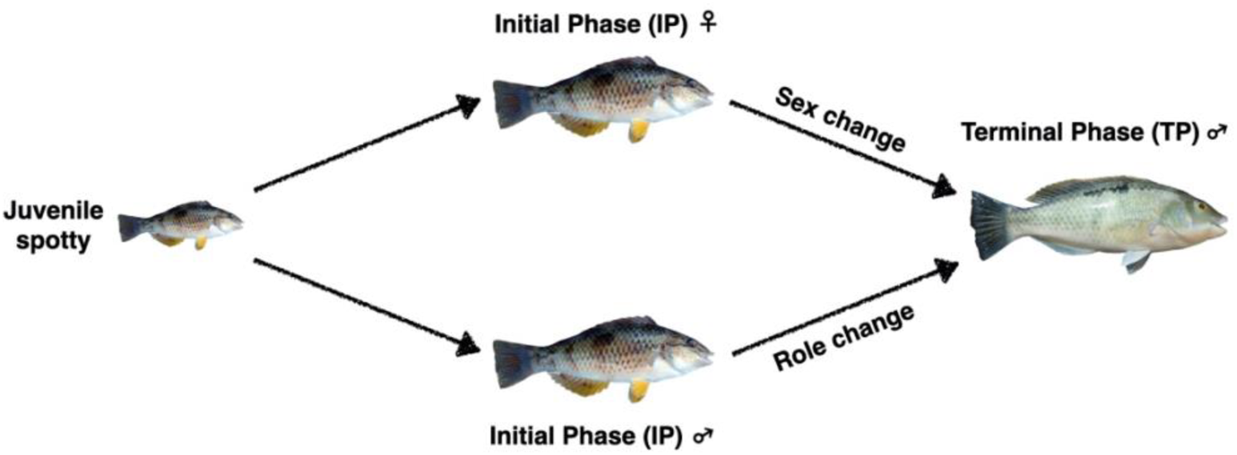
Life cycle of New Zealand spotty wrasse (*Notolabrus celidotus*). Juveniles first develop into either initial phase (IP) females or males, which can then become terminal phase (TP) males via sex or role change, respectively. Adapted from (Thomas et al., 2019; Todd et al., 2018). IP spotty wrasse image by Allan Burgess, TP spotty wrasse image by Jodi Thomas.

Sex change is effected through the reproductive axis, yet the underlying regulatory mechanisms are not well understood. The feminising and masculinising effects of the sex steroids, 17β-estradiol (E2) and 11-ketotestosterone (11KT) on sex changing fish are clear (Frisch, 2004; Kroon and Liley, 2000; Todd et al., 2016). However, the molecular interplay modulating their balance is complicated. Recent studies indicate that the glucocorticoid stress hormone cortisol can influence sex change. This is mediated via cross-talk between the hypothalamic-pituitary-interrenal (HPI) and hypothalamic-pituitary-gonadal (HPG) axes and involves the enzymes Cyp11c1 and Hsd11b2 (Liu et al., 2017; Todd et al., 2019). Studies in the tropical model bluehead wrasse also suggest involvement of the glucocorticoid (*nr3c1*) and mineralocorticoid (*nr3c2*) receptors (Todd et al., 2019). Indeed, alongside a direct effect of cortisol on the gonad through activation of the glucocorticoid receptors, the aforementioned adrenal enzymes could potentially modulate the estrogen-androgen balance, which remains to be fully elucidated. Comparing the interplay of these markers across the HPI and HPG axes of different model species will be important for clarifying the mechanistic basis of sex change.

The identification of an early molecular event that may trigger sex change is of special interest. The gonadal aromatase enzyme, responsible for the bioconversion of testosterone into E2 and encoded by *cyp19a1a,* is a strong candidate. An early decrease in E2 concentration has long been associated with the onset of sex change in several protogynous species (Bhandari et al., 2003; Liu et al., 2017; Muncaster et al., 2013; Nakamura et al., 1989). This is further supported by manipulative experiments where chemical inhibition of aromatase readily induces masculinisation of females in various species (Higa et al., 2003; Kroon et al., 2005; Lee et al., 2001; Nakamura et al., 2015; Nozu et al., 2009). The peptide anti-müllerian hormone (Amh) is implicated in regulating spermatogonial proliferation and male sex differentiation in fish (Li et al., 2015; Pfennig et al., 2015; Skaar et al., 2011; Zhang et al., 2020), and has also been identified as a possible early mediator of female-male sex change (Hu et al., 2015; Liu et al., 2017; Todd et al., 2019). With a network of candidate genes likely to influence the gonadal sex steroid environment, a targeted approach to characterise their expression across sex change is warranted.

In this study, we present histological, endocrine and gene expression data to characterise the sex change process in the spotty wrasse and establish this species as a new temperate model for the study of vertebrate sex change. We investigate molecular and endocrine pathways as well as potential triggers that may regulate gonadal restructure in these fish using both chemical and socially induced sex change.

## Materials and Methods

### Experimental set-up

#### Experiment 1: Induction of sex change in spotty wrasses by aromatase inhibition (AI2014)

In this experiment, the aromatase inhibitor (AI) fadrozole (C_14_H_13_N_3_) was used to induce sex change in captive IP spotty wrasse individuals between August and September 2014. Fish were captured around high tide by hook and line off the coast of Tauranga, Bay of Plenty, New Zealand (37.6878° S, 176.1651° E) and subsequently maintained at the Aquaculture Centre at Toi Ohomai Institute of Technology, Tauranga. TP males were distinguished from IP males (IPM) and females by external observation: IP fish have a large inky thumbprint spot in the middle of the body, whereas TP males have an irregularly shaped row of blackish spots and light electric blue wavy patterns on their cheeks (Choat, 1965) (Fig. 1). Thirty IP fish ranging from 154 – 229 mm total length (TL) were distributed across three 400-litre recirculating seawater systems under a natural photoperiod. Natural sex change was blocked by placing a TP male in each tank, creating a socially inhibitory environment. During the experiment, fish were fed frozen greenshell mussels (*Perna canaliculus*) three times per week. Pellets containing 200 μg fadrozole (Sigma-Aldrich) in a matrix of cholesterol:cellulose = 95:5 were made in-house (as described in (Lokman et al., 2015; Sherwood et al., 1988)). Sham pellets (vehicle) contained matrix only. Following an acclimation period of three weeks, on day 0 of the experiment all IP individuals were given a single intramuscular fadrozole implant (n=16) or a sham implant without hormone (n=14) using a Ralgun implanter (Syndel, Ferndale, WA).

All fish were removed from individual tanks on day 21 (Tank 1, n=11), day 39 (Tank 2, n=11) or day 60 (Tank 3, n=11; end of experiment). Fish were anesthetised in an aerated bath containing 600 ppm 2-phenoxyethanol (Sigma-Aldrich) and blood samples were collected from the caudal vein using a 1 mL heparinised syringe. Fish were then euthanised by rapid decapitation. The mid-section of one gonad was fixed in Bouin’s solution (TP males) or 10% neutral buffered formalin (IP individuals) overnight, and then stored in 70% EtOH at room temperature until paraffin embedding and sectioning for histological analysis. The same location in the gonads was used in all fish when fixing gonadal samples for histology. Body weight and length, and gonadal weight were measured for each fish.

#### Experiment 2: Social induction of sex change in spotty wrasses within their breeding season (SI2016)

Sex change was induced in captive spotty wrasses through manipulation of social groups (i.e., removal of males from the treatment tanks) between September and December 2016. Fifty IP and ten TP individuals ranging from 150 – 215 mm TL were captured and maintained as described in Experiment 1 (AI2014). Fish were distributed into groups across ten 400-litre recirculating seawater systems such that each tank contained a hierarchy of different sized IP fish and a single TP male (215 – 244 mm TL). After a 3-week acclimation, TP males were either removed from the treatment tanks (n = 5) or left in the control tanks (n=5). Subsequently, the largest IP fish from each tank was terminally sampled on days 0, 30, 50, 60, 65, or 66 (end of experiment). Blood plasma collection, anaesthetic administration, tissue dissection and recording of morphometrics were conducted as described for AI2014.

#### Experiment 3: Social induction of sex change in spotty wrasses outside their breeding season (SI2018)

Social manipulation was used to induce sex change in captive spotty wrasses outside the breeding season between January and April 2018. Sixty-five IP and twelve TP individuals ranging from 138 – 218 mm TL were captured and maintained as described in AI2014. Fish were distributed across twelve 400-litre recirculating seawater systems such that each tank contained a hierarchy of 4 – 5 different-sized IP fish and a single TP male (194 – 220 mm TL). Three control (5 IP females + 1 TP) tanks were maintained and nine manipulated (5 IP females - 1 TP) tanks had the males removed on day 0 after a 2-week acclimation period. A further five IP females were terminally sampled on day 0 to provide a baseline indication of reproductive status. Fish were sampled over a time series as follows; day 1 (n=5), day 11 (n=5), day 26 (n=10), day 36 (n=10), day 55 (n=10), day 92 (n=9). Eleven mortalities occurred during the experiment.

Anaesthetic administration, blood plasma collection, and recording of morphometrics were conducted as described for AI2014. One gonad was flash frozen in ice-cold (on dry ice) isopentane (C_5_H_12_) (Sigma-Aldrich) and stored at −80 °C for RNA analyses. The second gonad was preserved for histological analysis as described in AI2014.

Fish in all three experiments were maintained and manipulated in accordance with New Zealand National Animal Ethics Advisory Committee guidelines. Ethics applications were approved by the Animal Ethics Committee of Toi Ohomai Institute of Technology and independently reviewed by the University of Otago Animal Ethics Committee.

### Gonadal tissue processing for histology

Histological analysis of gonadal tissues was used to characterise cellular changes occurring across sex change. Tissues from AI2014 and SI2016, fixed in Bouin’s solution (TP males) or 10% neutral buffered formalin (IP individuals), were processed for routine embedding in paraffin (New Zealand Veterinary Pathology, Hamilton Laboratory, New Zealand). SI2018 gonadal tissues were embedded in paraffin (Otago Histology Services Unit, Department of Pathology, Dunedin School of Medicine, University of Otago, New Zealand). Sections were cut at 3 – 4 µm and stained with Mayer’s haematoxylin and eosin.

Gonadal sections were examined under light microscope to confirm the sex of each individual and samples were subsequently classified into a series of stages to demonstrate the transition from initial stage female through to terminal phase male (Table 1).

**Table 1.**

| Stage | Abbreviation | Characteristics |
| --- | --- | --- |
| **Initial Phase Female** | F | Ovary is dominated by healthy oocytes. Depending on season, these may be predominantly previtellogenic stage oocytes or may also include vitellogenic and maturing oocytes. Atretic late-stage oocytes may be common post-spawning. |
| **Early Transitioning** | ET | Atretic oocytes of all stages, including previtellogenic oocytes, common. Nests of gonial cells, yellow-brown bodies and stromal cells often evident. |
| **Mid-Transitioning** | MT | Spermatogenic cysts containing mainly spermatogonia or spermatocytes evident. Many atretic oocytes often still present. Stromal cells and cellular debris common. |
| **Late Transitioning** | LT | Evidence of lobular structure forming. Male germ cells dominate. Yellow-brown bodies and cellular debris common. |
| **Terminal Phase Male** | TP | Fully structured testis with spermatogenic cysts arranged into lobules. Sperm ducts developed peripherally inside the tunica albuginea and presence of central remnant lumen. All stages of spermatogenic germ cells may be present depending on season. A few degenerating oocytes may persist. |
| **Initial Phase Male** | IPM | Presence of lobule formation containing spermatogenic cysts. No evidence of degenerating oocytes. No central lumen and sperm ducts tend to be aggregated centrally. |

### Steroid measurements

Blood was centrifuged at 13,500 rpm for 3 minutes to obtain plasma, which was stored at −20 °C until steroid analysis. Measurement of blood plasma concentrations of E2 and 11KT across sex change were conducted by radioimmunoassay (RIA) after routine steroid extraction following procedures described in (Kagawa et al., 1981; Kagawa et al., 1982; Young et al., 1983). Assays were validated for spotty wrasse plasma, serially diluted plasma behaving in a manner similar to the standard curve (parallel displacement). Tritiated 11KT was synthesised using the methodology described in Lokman et al. (1997), whereas label for the E2 assay was acquired from Perkin Elmer. Antiserum against E2 was purchased from MyBioSource and antiserum against 11KT was kindly donated by Emeritus Professor Yoshitaka Nagahama, National Institute for Basic Biology, Okazaki, Japan. After incubation and separation of antibody-bound and -unbound steroid by charcoal-dextran solution (0.5% dextran/charcoal), tubes were centrifuged (15 min, 2000g), the supernatant was decanted, and radioactivity measured using a MicroBeta^®^ Trilux scintillation counter (Wallac 1450, Perkin Elmer). Samples from each experiment were run in separate assays with a minimum detectable level of 40 pg/tube (0.08 ng/mL) (E2) and 50 pg/tube (0.10 ng/mL) (11KT) for AI2014, 35 pg/tube (0.07 ng/mL) (E2) and 120 pg/tube (0.24 ng/mL) (11KT) for SI2016, and 50 pg/tube (0.10 ng/mL) (E2) and 70 pg/tube (0.14 ng/mL) (11KT) for SI2018. Extraction efficiencies were 46% (E2) and 82% (11KT) for AI2014; 61% (E2) and 87% (11KT) for SI2016; and 69% (E2) and 40% (11KT) for SI2018.

Due to non-normality of the RIA data, the non-parametric Kruskal–Wallis test (Kruskal and Wallis, 1952) was used to compare plasma steroid concentrations between sexual stages, for E2 or 11KT separately. If stage was found to have a significant effect, *post hoc* comparisons using Dunn’s tests (Dunn, 1961) with Benjamini–Hochberg correction for multiple comparisons (Benjamini and Hochberg, 1995) were performed between stages to determine where the significance lay, carried out in R (v. 1.1.453) (Core Team, 2013).

### RNA extraction from gonadal tissues

Gonadal tissues from the SI2018 experiment were used to assess sex change-related gene expression. Samples were homogenised using a power homogeniser before RNA extraction. RNA was extracted with Direct-zol RNA MiniPrep Plus (Zymo Research) without phase separation (on column DNase treatment).

RNA concentration was measured using a Qubit 2.0 Fluorometer (Life Technologies) and RNA integrity was evaluated on a Fragment Analyzer (Advanced Analytical Technologies Inc.). The RNA profiles of sex-changing gonads presented a strong peak of low molecular weight RNA. This is considered to be a result of massive 5S rRNA amplification in ovaries (Liu, 2016), and masks the 18S and 28S rRNA peaks used to calculate RNA Integrity Number (RIN) values, making them unreliable estimates of RNA integrity. Similar patterns have been observed in ovaries and/or intersex gonads of thicklip gray mullet (*Chelon labrosus* (Diaz de Cerio et al., 2012)), sharsnout seabream (*Diplodus puntazzo* (Manousaki et al., 2014)) and bluehead wrasse (Liu, 2016). Therefore, in spotty wrasses, RNA integrity for such samples was confirmed by visual inspection of 18S and 28S rRNA peaks.

### Gene expression analysis

A probe array of 18 candidate genes was designed for spotty wrasse based on the nanoString nCounter™ technology (Table S1). Spotty wrasse-specific transcript sequences were identified from a draft spotty wrasse transcriptome assembly (EV Todd & NJ Gemmell, 2015, unpublished data, now superseded by a fully annotated genome https://vgp.github.io/genomeark/Notolabrus_celidotus/). Reference transcripts from zebrafish (*Danio rerio*) were downloaded from GenBank or Ensembl for each target gene, and *actb1*, *eef1a1a* and *g6pd* as potential housekeeping genes (see Supplemental Materials for details on the determination of the housekeeping genes’ stability). Transcripts were used to identify the corresponding spotty wrasse sequences from the transcriptome via local BLASTn and tBLASTx (translated the query nucleotide sequences, and the spotty wrasse transcriptome into deduced amino acid sequences in all six possible frames, which were then compared by local alignment). For the best matching spotty wrasse contig, gene identity was confirmed using online nucleotide BLAST (BLASTn) against the NCBI database (http://www.ncbi.nlm.nih.gov/).

Best-match spotty wrasse contig sequences were submitted to nanoString Technologies for probe design. NanoString nCounter™ CodeSet gene expression quantification was performed on a panel of 18 sex change-related genes at the Otago Genomics Facility, Biochemistry Department, University of Otago, New Zealand. These data were generated using 100 ng gonadal RNA from the SI2018 fish (5 control females (day 0; F), 19 ET, 9 MT, 9 LT and 5 TP). The full suite of 18 genes were subjected to PCA analysis (see below) to help verify the gonadal staging. Here, we also present gene expression profiles for six gonadal genes relating to the endocrine function of the reproductive and stress axes (Table 2). The remaining expression data, including more novel targets, will be addressed in future work. The geometric mean of gene pair *actb1*|*g6pd* was selected as the reference gene for data normalisation (see Results). Normalisation calculations were performed automatically in the nanoString nSolver Analysis software (version 4.0), and normalised data were log base-2 transformed prior to analysis.

**Table 2.**

| Gene symbol | Gene description | Contig ID | Reference transcript ID |
| --- | --- | --- | --- |
| Housekeeping genes |  |  |  |
| *actb1* | β-actin, cytoplasmic 1 | c58053_g1_i1 | NM_131031.1 |
| *eef1a1a* | eukaryotic translation elongation factor 1 alpha 1a | c58053_g1_i1 | NM_200009.2 |
| *g6pd* | glucose-6-phosphate dehydrogenase | c39960_g1_i1 | ENSDART00000104834.6 |
| Steroidogenic enzymes and hormone receptors |  |  |  |
| *cyp19a1a* | aromatase a (gonad isoform) | c52027_g1_i1 | NM_131154.3 |
| *cyp11c1/b2* | steroid 11β-hydroxylase | c62027_g1_i1 | NM_001080204.1 |
| *hsd11b2* | 11β-hydroxysteroid dehydrogenase type 2 | c67035_g1_i1 | NM_212720.2 |
| *nr3c1* | glucocorticoid receptor | c36910_g2_i1 | NM_001020711.3 |
| *nr3c2* | mineralocorticoid receptor | c49976_g1_i2 | NM_001100403.1 |
| Key sex-related transcription factors |  |  |  |
| *amh* | anti-Müllerian hormone | c51546_g1_i1 | NM_001007779.1 |

Due to non-normality of the normalised nanoString data, the non-parametric Kruskal–Wallis test (Kruskal and Wallis, 1952) was run separately for each target gene to determine whether stage had a significant effect on gene expression level, and *post hoc* comparisons using Dunn’s tests (Dunn, 1961) with Benjamini–Hochberg correction for multiple comparisons (Benjamini and Hochberg, 1995) were performed to determine where the significance, if any, lay between stages. All analyses were performed in R (v. 3.3.2) (Core Team, 2013). NanoString Expression Data Analysis Guidelines (MAN-C0011-04) were followed to determine an expression threshold, set as the log_2_ of the geometric mean of the negative control counts plus two standard deviations.

Principal Component Analysis (PCA) (scaled, R (v. 1.1.453) (Core Team, 2013)) was used to visualise variation in the overall mRNA expression across the expanded suite of 18 sex change-related genes (Table S1) in relation to sexual stage and assess clustering of the histological defined in Table 1. To identify the genes contributing most to observed patterns represented by the first two principal components, component loadings (defined as eigenvectors scaled by the square root of the respective eigenvalues) were represented as coordinates in a Cartesian plane.

## Results and Discussion

The definition and delineation of sexual stages differs between studies of protogynous fish. In this study, we define five gonadal stages to describe the sexual transition from female to terminal phase male endpoints in the temperate spotty wrasse. While these stages are based on gonadal histology the classification is also supported by the PCA analysis of the gonadal gene expression data. These samples showed clusters arranged in the sequential order of the sexual stages from female through to male (PC1, 50.7% variation explained; Fig. 2).

**Figure 2.**
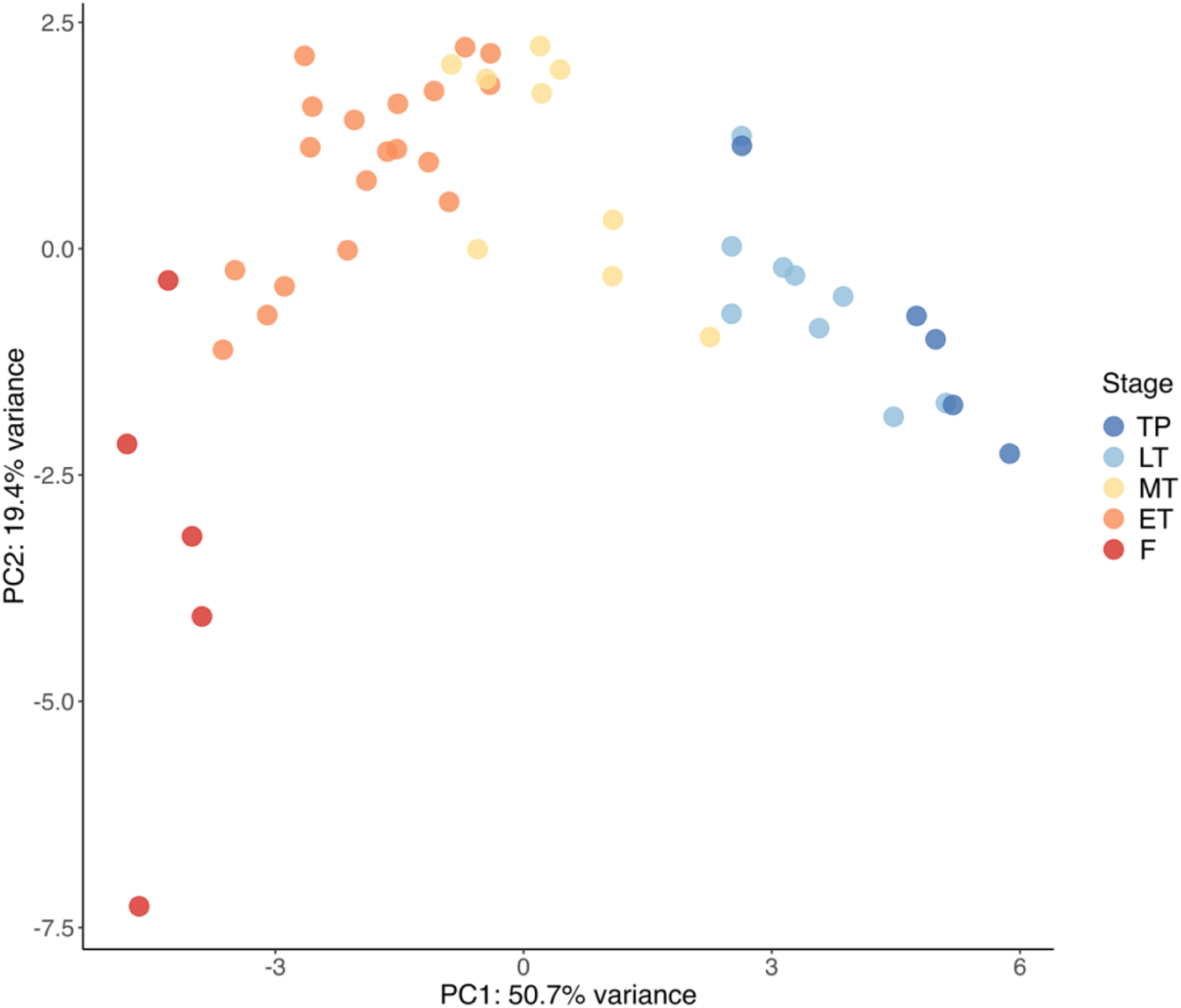
PCA (18 genes) of gonad samples. The transition from females to males is captured along PC1 (50.7% variance), whereas PC2 (19.4% variance) extremes represent sexually differentiated gonads (bottom) and undifferentiated transitionary gonads (top). Sample sizes: F n = 5, ET n = 19, MT n = 9, LT n = 9, TP n = 5. Abbreviations: control female (F), early transitioning fish (ET), late transitioning fish (LT), mid-transitioning fish (MT), terminal phase male (TP).

The spotty wrasse gonadal gene set was also clustered according to the degree of their developmental commitment (PC2, 19.4% variation explained). Fully differentiated ovary and testis were separated from less differentiated transitional gonads. These patterns show a striking resemblance to whole transcriptome gonadal expression profiles from transitioning tropical bluehead wrasse (Todd et al., 2019). This demonstrates the importance and relevance of the spotty wrasse gene panel to studying sexual transition in protogynous wrasses more broadly.

### Histological analysis of gonadal sex change

#### Experiment 1: Induction of sex change in spotty wrasses by aromatase inhibition (AI2014)

Aromatase inhibition successfully induced sex change in 93% of the surviving (2 mortalities) captive female spotty wrasses held under socially inhibitory conditions. Histological analysis confirmed that among the AI-implanted females, 12 fish reached ET stage (day 21, n=2; day 39, n=5; day 60 n=5), and one reached LT stage (day 60). A single fadrozole-implanted fish remained female. In contrast, none of the control females showed signs of ovarian atresia or sex change. None of the experimental fish were found to be IPM.

#### Experiment 2: Social induction of sex change in spotty wrasses within their breeding season (SI2016)

The manipulation of social groups, through male removal, successfully promoted sex change (81%) in female spotty wrasses during the 2016 breeding season. Histology confirmed that among the socially manipulated females, 15 fish reached ET stage (day 30, n=2; day 50, n=3; day 60, n=4; day 65, n=3; day 66, n=3), one reached MT stage (day 50), one LT stage (day 50), and one was found to be a fully TP male (day 60). Four of the socially manipulated fish remained female (day 30, n=2; day 66, n=2). No conclusive evidence of sex change was found in any of the control females, although ovarian atresia was observed in four of these individuals (day 30, n=3; day 66, n=1). Histological analysis also confirmed that five of the starting 50 IP individuals were IPM (10.0% frequency), a slightly higher ratio than reported previously (4.1 – 5.7%) (Jones, 1980).

#### Experiment 3: Social induction of sex change in spotty wrasses outside their breeding season (SI2018)

Male removal conducted outside the breeding season also induced sex change in female spotty wrasses. All socially manipulated females showed histological signs of ovarian degeneration or sex change. Histology confirmed that 18 females reached ET stage (day 1, n=3; day 11, n=2; day 26, n=3; day 36, n=3; day 55, n=4; day 96, n=3), 12 MT stage (day 1, n=1; day 11, n=3; day 26, n=4; day 36, n=1; day 55, n=3), 13 LT stage (day 26, n=2; day 36, n=6; day 55, n=3; day 92, n=2) and 6 became full TP males (day 1, n=1; day 92, n=5). No IPM were found (Fig. 3). Unfortunately control tanks with male fish present experienced an unknown infection and reduced survival in the later stages of the experiment. This confounded the efficacy of socially induced sex change in this experiment. Nonetheless, the results of the other two experiments (AI2014 & SI2016) indicate that both chemical and social manipulation increases sex change compared to control female fish that have a male present (Fisher’s exact test, p < 0.001). This is also supported by previous studies with this species (Muncaster, unpublished data).

**Figure 3.**
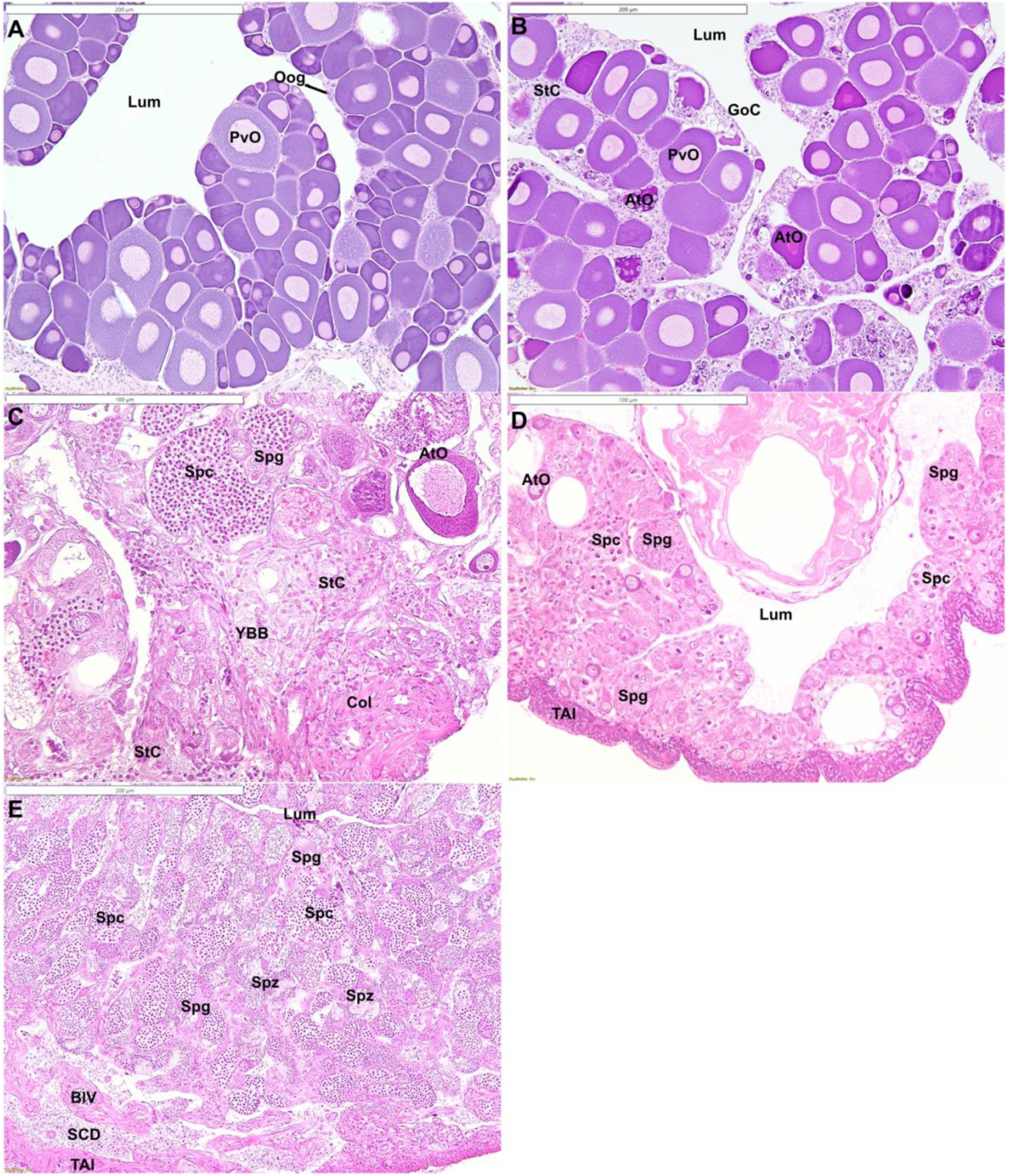
Histological stages of gonadal development in spotty wrasse: A) female; B) early transitioning fish; C) mid-transitioning fish; D) late transitioning fish; E) terminal phase male. Abbreviations: atretic oocyte (AtO), blood vessel (BlV), collagen (Col), gonial cells (GoC), gonadal lumen (Lum), previtellogenic oocyte (PvO), oogonia (Oog), stromal cells (StC), sperm collecting duct (SCD), spermatocytes (Spc), spermatogonia (Spg), spermatozoa (Spz), tunica albuginea (TAl), yellow-brown body (YBB). Scale bars in A, B, E = 200 µm; C, D = 100 µm.

### Incidence of sex change depending on seasonality

The breeding season of spotty wrasses in northern New Zealand lasts from late July until the end of November and peaks in the austral spring (Jones, 1980). Of the socially manipulated spotty wrasses, the more mid-transitional and fully transformed males were recorded in the experiment conducted during the post-spawning period (SI2018) in summer and early autumn (January to April; Fig. 4). In this experiment, 55% of IP fish had MT through to TP stage gonads, whereas less than 25% of fish socially manipulated during the October-December breeding season (SI2016) presented the same stage gonads. Sex change is often seasonally biased with the greatest occurrence following the breeding season in temperate and warm water species, but occurs year-round in many tropical species (Alonso-Fernández et al., 2011; Li et al., 2007; Muncaster et al., 2013; Sadovy and Shapiro, 1987; Thomas et al., 2019). Our results support observations in wild spotty wrasses, in which natural sex change has been documented during the non-reproductive months between November and May (Jones, 1980). This indicates that for experimental purposes, post-spawned fish present the best candidates for socially manipulated sex change. However, we have also demonstrated that male removal can lead to sex change within the breeding season.

**Figure 4.**
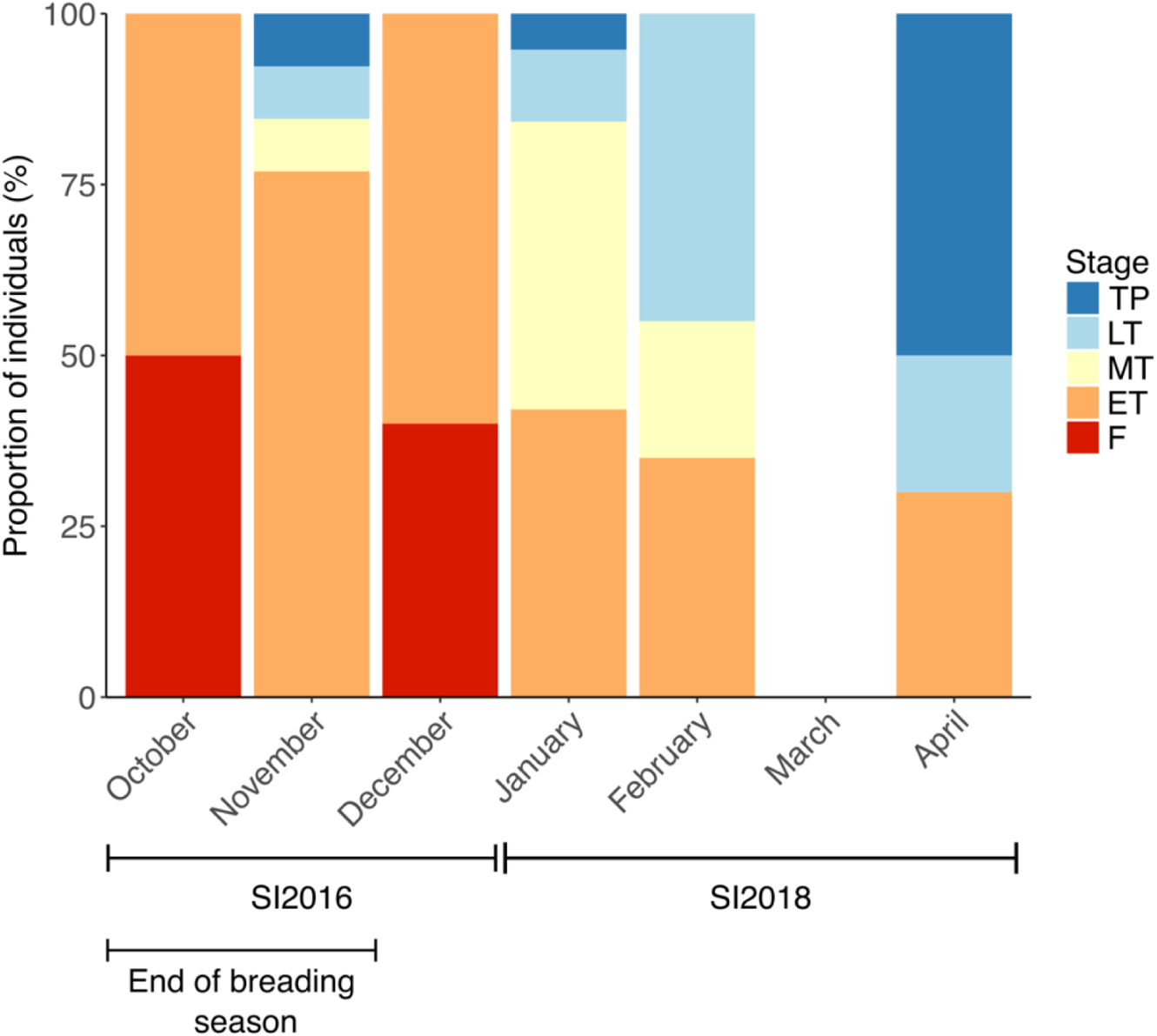
Proportion of SI2016 and SI2018 spotty wrasses in different sexual phases depending on month of sampling. Abbreviations: initial phase female (F), early transitioning fish (ET), late transitioning fish (LT), mid-transitioning fish (MT), social induction experiment 2016 (SI2016), social induction experiment 2018 (SI2018), terminal phase male (TP). Sample sizes: October n=4, November n=13, December n=5, January n=20, February n=20, March n=0, April n=9.

### Steroid production in relation to sex change

Plasma E2 concentrations showed a general decreasing trend from female to male stages (Fig. 5A). Fish treated with an aromatase inhibitor (AI2014) had mean plasma E2 concentrations ranging from 0.08 ng/ml to 0.34 ± 0.25 SD ng/ml for LT and ET stages respectively. In comparison, female fish had a higher mean plasma E2 concentration of 0.42 ± 0.46 SD ng/ml, although this was not statistically different. Social manipulation during the breeding season (SI2016) showed significantly higher plasma E2 concentrations in females (0.38 ± 0.39 SD ng/mL) compared to males (0.07 ± 0.00 SD ng/mL, p < 0.001 for TP and 0.07 ± 0.00 SD ng/mL, p < 0.05 for IPM). However, plasma E2 concentrations were minimal in all sexual stages when social manipulation was conducted after the completion of the spawning season (SI2018). Similar reduced E2 concentrations after breeding were observed in other protogynous species (Bhandari et al., 2003; Li et al., 2007; Muncaster et al., 2010). Gonadal resorption and quiescence often follows reproduction in temperate fish and this is characterised by ovarian atresia and reduced sex steroid concentrations (Scott et al., 1984). This is not surprising considering the importance of E2 in driving seasonal oocyte growth in teleosts (Jalabert, 2005; Lubzens et al., 2010; Nagahama, 1994; Patiño and Sullivan, 2002).

**Figure 5.**
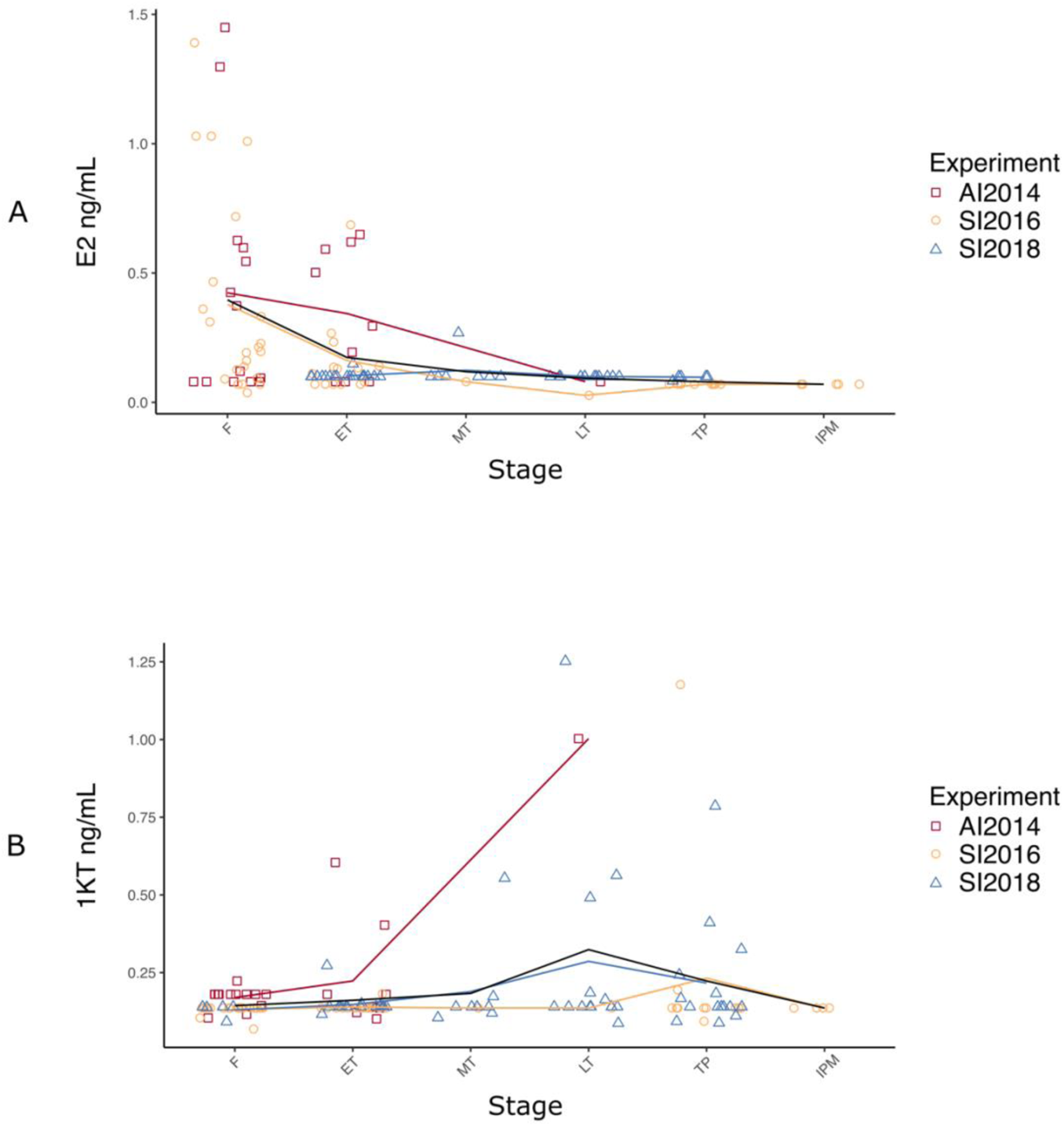
Plasma E2 (A) and 11KT (B) concentrations in females, early, mid- and late transitioning fish, and TP and IP males obtained across the three experiments, AI2014, SI2016 and SI2018. Each red (AI2014), yellow (SI2016) and blue (SI2018) line represents the variation in mean E2 or 11KT concentrations across groups per experiment while the black line represents the variation in overall mean concentrations for the three experiments altogether. Sample sizes: E2, F n = 36, ET n = 42, MT n = 8, LT n = 14, TP^☨^ n = 17, IP n = 5; 11KT, F n = 40, ET n = 41, MT n = 9, LT n = 15, TP^☨^ n = 26, IP n = 4. Abbreviations: 11-ketestosterone (11KT), aromatase inhibition experiment 2014 (AI2014), initial phase female (F), 17β-estradiol (E2), early transitioning fish (ET), initial phase male (IPM), late transitioning fish (LT), mid-transitioning fish (MT), social induction experiment 2016 (SI2016), social induction experiment 2018 (SI2018), terminal phase male (TP). ^☨^Both control males used throughout the experiments to create a socially inhibitory environment (SI2016: E2 n = 10, 11KT n = 10; SI2018: E2 n = 0, 11KT n = 10), and males obtained through sex change of manipulated females (SI2016: E2 n = 1, 11KT n = 1; SI2018: E2 n = 5, 11KT n = 5) were grouped altogether as TP males for the purpose of this analysis.

While E2 has a clear role in maintaining ovarian function, there is no obvious relationship between plasma E2 concentrations and the initiation of sex change in spotty wrasses. A marked decrease in E2 concentrations has been implicated as a critical initiator of sex change in many protogynous species (Bhandari et al., 2003; Liu et al., 2017; Muncaster et al., 2013; Nakamura et al., 1989). Yet, despite a 2.3-fold decrease, there was no significant difference in plasma E2 concentration between F and ET (0.16 ± 0.16 SD ng/mL) fish during the breeding season (SI2016). Similarly, there was no significant decrease of plasma E2 concentration between female and early transitional spotty wrasses from the other experiments (AI2014 & SI2018). While this may, in part, relate to sample size or seasonal influence, similar results were also reported in the bambooleaf wrasse (*Pseudolabrus sieboldi*) (Ohta et al., 2008) and orange-spotted grouper (*Epinephelus coioides*) (Chen et al., 2020b). Undetectable E2 concentrations existed from MT to male (TP & IP) stages in nearly all fish. Therefore, while there is no evidence of a minimum plasma E2 threshold required to initiate gonadal transition in spotty wrasse, a general reduction is associated with the process of sex change in this species. Indeed, E2 depletion leads to masculinisation in hermaphroditic and gonochoristic species alike (Bhandari et al., 2004; Li et al., 2019; Nozu et al., 2009; Paul-Prasanth et al., 2013; Takatsu et al., 2013).

Elevated plasma 11KT concentrations were observed in individual fish towards the transitional and male stages in all three experiments (AI2014, SI2016 and SI2018; Fig. 5B). While variability in the data and reduced statistical power made the detection of discernible differences impossible, the timing of these observations coincides with the histological appearance of spermatogenic cysts (see Table 1). Increased 11KT concentrations were also evident from mid transition onwards in other protogynous species (Bhandari et al., 2003; Nakamura et al., 1989; Nakamura et al., 2005). Fish treated with AI (AI2014) had minimal 11KT values in the earlier sexual stages. This was evident in the F (0.17 ± 0.03 ng/ml) and ET (0.22 ± 0.17 SD ng/ml) fish while a later stage LT individual presented with 1.0 ng/ml. Fish socially manipulated during the breeding season (SI2016) showed remarkably similar plasma 11KT concentrations regardless of sexual stage. These values ranged from 0.13 ± 0.02 SD ng/ml in F to 0.23 ± 0.31 SD ng/ml in TP. All of the F, transitional, and IPM had either identical mean or individual 11KT concentrations of 0.14 ng/ml. Many of the androgen concentrations of fish socially manipulated after the breeding season (SI2018) remained minimal as expected during quiescence.

The lowest mean 11KT concentrations (0.13 ± 0.02 SD ng/ml) presented in F and were only slightly higher in subsequent stages such as ET (0.15 ± 0.03 SD ng/ml) and MT fish (0.19 ± 0.15 SD ng/ml). Highest plasma 11KT values were observed in LT (0.29 ± 0.32 SD ng/ml) fish, while slightly reduced concentrations were recorded in TP individuals (0.22 ± 0.18 SD ng/ml). This disparity of androgen concentrations in late-stage fish most likely reflects the chronology of the experiment. TP fish were removed from the socially manipulated tanks at the beginning of the experiment immediately after the breeding season. Collection of these tissue and plasma samples would likely have been when spermatogenesis was minimal or non-existent. The occurrence of several LT fish occurred later in the experiment, coinciding with seasonal gonadal recrudescence.

The androgen profiles from the fish in this study do not show a clear statistical relationship between sexual stages. However, the role of 11KT in driving spermatogenesis is well established in teleosts (Miura et al., 1991; Nagahama, 1994; Nakamura et al., 1989; Schulz et al., 2010). Much of this androgen activity is likely to be paracrine in nature with steroidogenic somatic cells stimulating local germ cells both directly and indirectly within the gonadal compartment (Schulz, 1986). The additional role of 11KT in expressing seasonal male secondary sexual characteristics, such as morphometric and behavioural modifications, also exists (Borg, 1994; Semsar and Godwin, 2004). This likely requires elevated plasma concentrations for remote, effective target cell signalling and may in part explain substantial elevations of 11KT prior to breeding in many male teleosts. The absolute concentrations of androgen required to stimulate spermatogenesis are conceivably much lower. Furthermore, prior to these peak physiological concentrations the actual concentrations of 11KT within the gonad are likely to be higher than in the plasma. It is, therefore, possible that the absolute concentrations of 11KT required to initiate spermatogenesis during gonadal restructure are not reflected in the spotty wrasse plasma samples. This issue may be further investigated using *in vitro* explant culture systems (Goikoetxea et al., 2020). Alternatively, gonadal expression of key steroidogenic enzymes should also reflect, in proxy, the androgen activity across different sexual stages.

### Sexually dimorphic expression of gonadal endocrine genes

To obtain a more sensitive assessment of the endocrine regulation of gonadal sex change, the expression of three genes encoding gene products involved in steroid biosynthesis, two steroid receptors, and anti-Mullerian hormone were measured. The temporal regulation of steroid hormones is essential for the coordination of sex differentiation, sexual maturation and behaviour in vertebrates. They are also potent mediators of gonadal sex change in teleosts (Guiguen et al., 2010; Higa et al., 2003; Nakamura et al., 1989). As expected *cyp19a1a* expression was greatest in F and was gradually downregulated across sex change (Fig. 6A). Expression did not differ significantly across stages. As reflected in the E2 data, there was no evidence of an early, rapid downregulation that has been thought to trigger sex change in other species (Gemmell et al., 2019; Liu et al., 2017; Todd et al., 2016). Levels of *cyp19a1a* expression remained similar across ET, MT and LT stages (2018) while in comparison, plasma E2 concentrations were negligible by mid transition in all three experiments (AI2014, SI2016 and SI2018). While *cyp19a1a* is an unlikely proximate trigger of sex change in spotty wrasse, a more distant association exists nonetheless. This is evident in the number of fish that changed sex following aromatase inhibition (AI2014) as well as the occurrence of sex change in fish socially manipulated after the breeding season (SI2018) when plasma E2 concentrations were minimal. Considering the potent feminising action of E2, a reduction of gonadal concentrations may act as a gateway to facilitate the progression of transition rather than acting as an early trigger. This action may be through the release of steroid-induced suppression on male-pathway genes (Guiguen et al., 2010).

**Figure 6.**
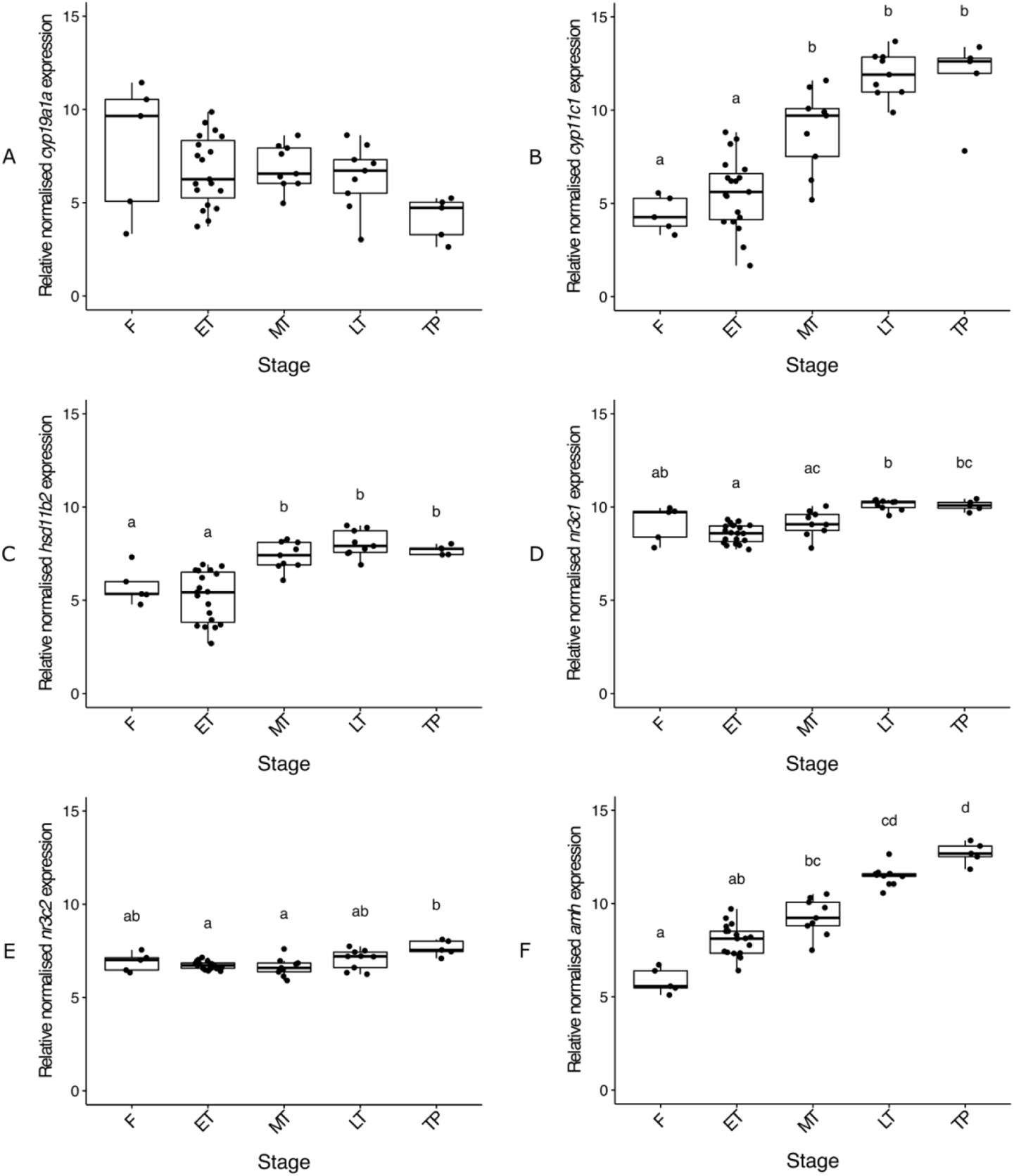
Relative gonadal expression of *cyp19a1a* (A), *cyp11c1* (B), *hsd11b2* (C), *nr3c1* (D), *nr3c2* (E) and *amh* (F) mRNA. Expression levels are compared among control females sampled on day 0, transitioning individuals and TP males. In the boxplots, each point represents an individual fish, the middle bold line represents the median, the edges of the box represent the upper and lower quartiles, and vertical lines represent the minimum and maximum values. Letters denote a significant difference in distribution between groups. Sample sizes: F n = 5, ET n = 19, MT n = 9, LT n = 9, TP^☨^ n = 5. Abbreviations: control female (F), early transitioning fish (ET), late transitioning fish (LT), mid-transitioning fish (MT), terminal phase male (TP). ^☨^Both a male used during the acclimation period of the experiment (n = 1), and males obtained through sex change of socially manipulated females (n = 4) were grouped altogether as TP males for the purpose of this analysis.

In teleosts, testosterone (T) can be bioconverted into 11KT by the enzymes 11β-hydroxylase (Cyp11c1) and 11β-hydroxysteroid dehydrogenase type 2 (Hsd11b2) (Frisch, 2004). Upregulation of *cyp11c1* was observed across sex change (Fig. 6B), with significantly (*X^2^* (4) = 32.39, p < 0.001) greater expression in TP spotty wrasses than F (median 2.6-fold greater) and ET (median 2.1-fold greater) fish. Albeit less pronounced, *hsd11b2* expression increased in a similar pattern across sex change (*X^2^* (4) = 32.13, p < 0.001; Fig. 6C). The simultaneous upregulation of both *cyp11c1* and *hsd11b2* from MT through to TP stages coincides with the presence of spermatogenic cysts in the gonad and likely drives the increase in gonadal 11KT production. Although this is not evident in plasma 11KT concentrations (see Fig. 5B), these expression patterns may indicate either undetected paracrine action of 11KT within the early testis or the accumulation of enzyme transcripts prior to translation.

Another contemporary hypothesis for a universal trigger of sex change involves stress, Cyp11c1 and Hsd11b2 are key players in the teleost stress response, being necessary for the interrenal biosynthesis of cortisol and for its subsequent inactivation to cortisone, respectively (Arterbery et al., 2010; Fernandino et al. 2013; Goikoetxea et al., 2017). Cross-talk between the interrenal and reproductive axes through the upregulation of these enzymes has been implicated as influencing masculinisation in teleosts (Chen et al. 2020; Fernandino et al, 2013; Goikoetxea, 2020; Hattori et al. 2013; Liu et al., 2017, Solomon-Lane et al. 2013), particularly through the production of weak interrenal-derived androgens, released as part of the stress response, which could conceivably serve as precursors for 11KT synthesis (Lokman et al., 2002). As cortisol exerts its effects by activating its receptors, among which Nr3c1 and Nr3c2 are best characterised, the genes encoding these receptors present interesting targets. These genes have opposing expression patterns in bluehead wrasse, with *nr3c1* showing a male bias while *nr3c2* is female-biased (Liu et al., 2015; Todd et al., 2019). However, gonadal expression of both *nr3c1* (*X^2^* (4) = 28.40, p < 0.001) and *nr3c2* (*X^2^* (4) = 13.02, p < 0.05) was elevated in the LT to TP stages of spotty wrasse (Figs. 6D and 6E).

Expression of these genes did not differ between F and TP fish, which is likely connected to sample size-related variability, particularly in female fish. Early-stage expression of *cyp11c1*, *hsd11b2* and *nr3c2* was thought to indicate a role for cortisol in triggering sex change in the bluehead wrasse model (Todd et al., 2019). In contrast, upregulation of these genes occurred in later stage gonads in the spotty wrasse. This suggests a relationship between cortisol and sex change may exist but not necessarily as an initiator in this species. Chen et al. (2020a) report a marked transient increase of cortisol concentrations during the early stages of protogynous sex change in orange-spotted grouper. Similar to what we observed in spotty wrasse, these fish do not show a significant reduction in levels of E2 during sex change. As such, the precise role of cortisol and the stress response during sex change warrants further investigation.

Anti-Müllerian hormone, encoded by *amh*, is strongly linked to male vertebrate sex differentiation (Josso et al., 2001; Pfennig et al., 2015). This was reflected in a linear function across sex change stages in spotty wrasse. *Amh* expression was greatest in TP males (2.2-fold, 1.6-fold and 1.4-fold greater in TP than F, ET and MT, respectively; *X^2^* (4) = 38.71, p < 0.001; Fig. 6F). The apparent upregulation of *amh* in ET and MT stage fish indicates its importance in the initiating stages of sex change in this species. Furthermore, *amh* demonstrated the greatest contribution to PC1 (ovary-to-testis transition) of the PCA analysis. This is consistent with other studies showing upregulation of *amh* at the onset of protogynous sex change and complementary downregulation in the early stages of protandrous sex change (Hu et al., 2015; Liu et al., 2017; Wu et al., 2015). The expression pattern of *amh* provides further validation for the histological staging of ET individuals as resolving cellular differences between ET fish and post-spawned females with regressing ovaries can be difficult in sex changing fish. A recent mechanistic model for socially induced sex change in the protogynous bluehead wrasse highlights an association between early *amh* activation and the stress hormone, cortisol (Todd et al., 2019). Therefore, *amh* may form part of the molecular trigger that initiates sex change, and its upregulation could be a useful early molecular marker for protogynous species.

## Conclusion

This study validates the chemical and social induction of sex change in captivity and describes the anatomical, endocrine and molecular events during sex change to establish the New Zealand spotty wrasse as a temperate-water model for sex change research. Sex change is most readily induced following the reproductive season when plasma sex steroid concentrations are low.

In general, the female-related markers E2 and *cyp19a1a* declined during sex change while the male-related markers 11KT, *cyp11c1*, *hsd11b2* increased. The enzymes encoded by these latter genes are also actively involved in the interrenal stress response. The gonadal expression of *nr3c1* and *nr3c2* increased in the later stages of sex change in spotty wrasse. This could suggest a putative role for stress during sex change but there is no clear evidence for cortisol as an early-stage activator of sex change. Similarly, E2 did not seem to be an initiator of sex change in this model, but *amh* showed early-stage upregulation. Apart from being an interesting target to better understand early activators of sex change, *amh* may prove to be a beneficial marker to resolve ET stage fish from females undergoing seasonal ovarian atresia

## Supporting information

Supplementary Methods Refs Figures Table

## Notes

### Competing Interest Statement

The authors have declared no competing interest.

