## Supplementary Methods Refs Figures Table for "Sex change in aquarium systems establishes the New Zealand spotty wrasse (*Notolabrus celidotus*) as a temperate model species for the investigation of sequential hermaphroditism"

**Supplementary Materials and Methods**

Analysis of stability of housekeeping genes:

Two approaches, RefFinder (https://www.heartcure.com.au/reffinder/?type=reference) (Xie et al. 2012) and BestKeeper (https://www.gene-quantification.de/bestkeeper.html) (Pfaffl et al. 2004) were used to determine the stability of gene expression of the potential housekeeping genes (*g6pd*, *eef1a1a* and *actb1*) and their suitability as reference genes for the normalisation of nanoString results. However, neither RefFinder or BestKeeper considers potential differences in reference gene expression between experimental groups, which can confound results and lead to an inaccurate interpretation (Dheda et al. 2004; Setiawan and Lokman 2010). Therefore, candidate reference genes were also evaluated for differences in target molecule counts between sex-change stages. Due to non-normality of the raw nanoString data, the non-parametric Kruskal–Wallis test (Kruskal and Wallis 1952) was used on the *actb1*, *eef1a1a* and *g6pd* counts, as well as the geometric mean of all possible combinations of these three genes, to determine whether stage had a significant effect on expression levels of each candidate housekeeping gene.

The ranking of candidate reference genes by RefFinder, Δ CT and NormFinder was *actb1* > *g6pd* > *eef1a1a*. The BestKeeper ranking was *g6pd* > *actb1* > *eef1a1a*, based on both SD and r (Tables S2A and S2B). These results suggest that *actb1* and *g6pd* should be used as reference genes for normalisation of the nanoString data.

Non parametric testing (Kruskal–Wallis; (Kruskal and Wallis 1952)) showed that gonadal expression of the candidate reference genes *actb1* and *eef1a1a* were significantly affected by sex-change stages (*actb1*, $X^2$ (4) = 23.94, $p < 0.001$; *eef1a1a*, $X^2$ (4) = 29.53, $p < 0.001$). The geometric mean of the mRNA levels of all three genes and that of gene pairs *actb1*|*eef1a1a* and *eef1a1a*|*g6pd* were also significantly influenced by stage (*actb1*|*eef1a1a*|*g6pd*, $X^2$ (4) = 16.77, $p < 0.005$; *actb1*|*eef1a1a*, $X^2$ (4) = 27.53, $p < 0.001$; *eef1a1a*|*g6pd*, $X^2$ (4) = 13.82, $p < 0.01$). Candidate reference gene *g6pd* mRNA levels ($X^2$ (4) = 7.97, p = 0.09) and the geometric mean of gene combination *actb1*|*g6pd* mRNA levels were not significantly affected by stage; ($X^2$ (4) = 9.03, p = 0.06) (Fig. S1). Consequently, gene pair *actb1*|*g6pd* was selected to normalise the target gene expression data (i.e., highest p-value for a gene combination observed) (Fig. S1E).

### Figure legends

Figure S1. Relative mRNA levels of each candidate reference gene, *actb1* (A), *eef1a1a* (B), *g6pd* (C); geometric mean of *actb1* and *eef1a1a* (D), geometric mean of *actb1* and *g6pd* (E), geometric mean of *eef1a1a* and *g6pd* (F), and geometric mean of *actb1*, *eef1a1a* and *g6pd* (G) in the gonad. *indicates a significant effect of sexual stage on relative mRNA levels. Sample sizes: F n = 5, ET n = 19, MT n = 9, LT n = 9, TP† n = 5. Abbreviations: control female (F), early transitioning fish (ET), late transitioning fish (LT), mid-transitioning fish (MT), terminal phase male (TP). †Both a male used during the acclimation period of the experiment (n = 1), and males obtained through sex change of socially manipulated females (n = 4) were grouped altogether as TP males for the purpose of this analysis.

Table S1. Genes analysed in spotty wrasse gonad using the nanoString nCounter™ CodeSet technology. Abbreviations: high mobility group (HMG), sex-determining region (SRY).

Table S2. Overview of potential reference genes from gonadal samples ranked from higher to lower stability (top to bottom) using different statistical approaches. RefFinder, Δ CT, BestKeeper SD and NormFinder rankings were obtained in RefFinder (A), while BestKeeper r was calculated on excel-based BestKeeper (B). Abbreviations: average (Ave.), geometric mean (GM), Pearson's correlation coefficient (r), standard deviation (SD), stability value (SV).

**Figures:**

Figure S1

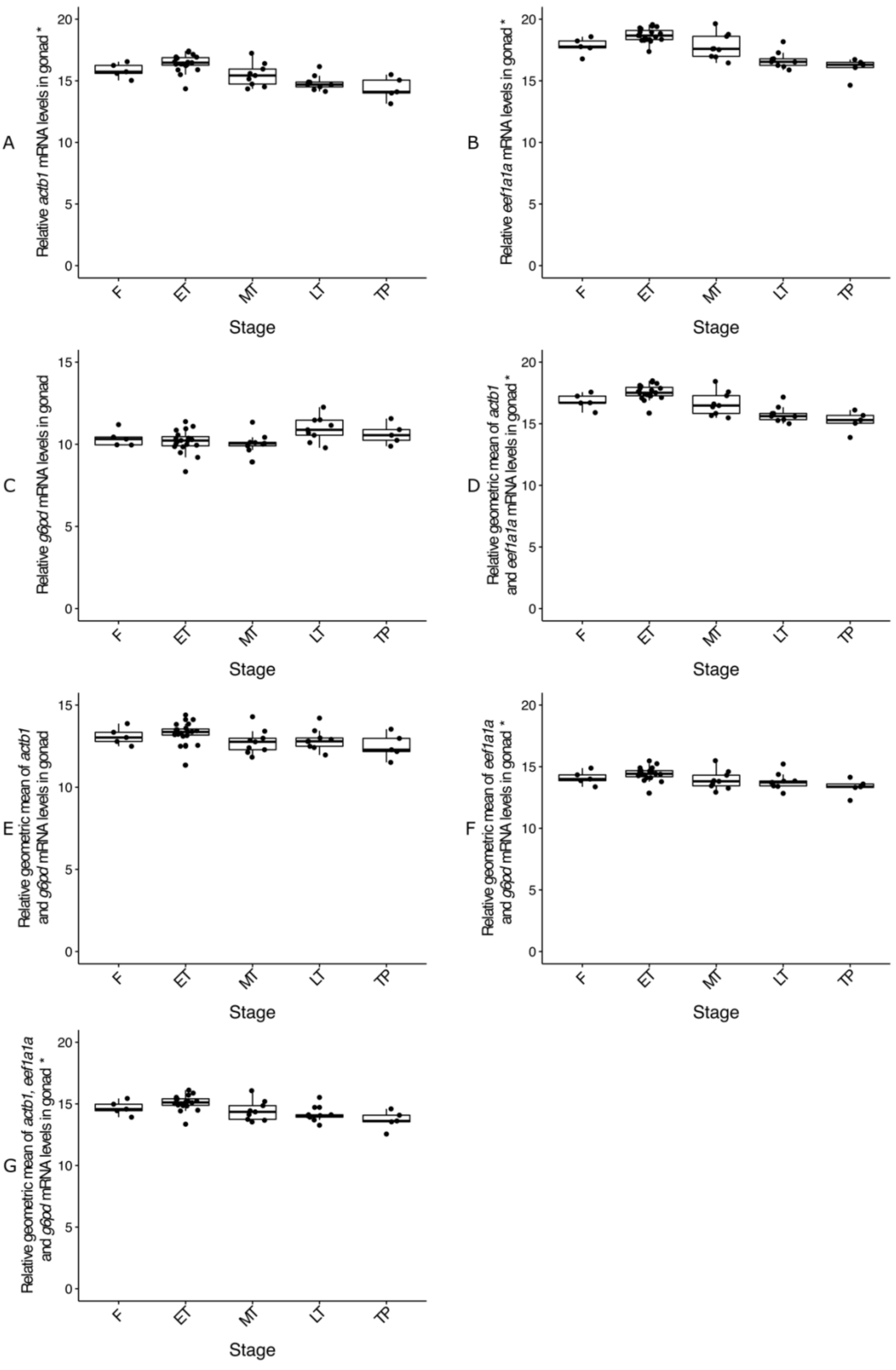

Table S1

| Gene symbol | Gene description | Contig ID | Reference transcript ID |
|---|---|---|---|
| Steroidogenesis and hormone receptors | | | |
| *cyp19a1a* | aromatase a (gonad isoform) | c52027_g1_i1 | NM_131154.3 |
| *cyp11c1/b2* | steroid 11ß-hydroxylase | c62027_g1_i1 | NM_001080204.1 |
| *hsd11b2* | 11ß-hydroxysteroid dehydrogenase type 2 | c67035_g1_i1 | NM_212720.2 |
| *nr3c1* | glucocorticoid receptor | c36910_g2_i1 | NM_001020711.3 |
| *nr3c2* | mineralocorticoid receptor | c49976_g1_i2 | NM_001100403.1 |
| Key sex-related transcription factors | | | |
| *foxl2a* | forkhead box L2a | c53356_g1_i1 | NM_001045252.2 |
| *dmrt1* | doublesex and mab-3 related transcription factor 1 | c66498_g1_i1 | NM_205628.2 |
| *amh* | anti-Müllerian hormone | c51546_g1_i1 | NM_001007779.1 |
| *sox9a* | SRY-related HMG box 9a | c53707_g2_i1 | NM_131643.1 |
| Rspo1/Wnt/ß-catenin pathway | | | |
| *ctnnb1* | catenin (cadherin-associated protein), beta 1 | c47984_g1_i2 | NM_131059.2 |
| *rspo1* | R-spondin-1 (precursor) | c50451_g1_i2 | NM_001002352.1 |
| E3 ubiquitin-protein ligase | | | |
| *znrf3* | zinc and ring finger 3 | c68386_g1_i1 | NM_001308555.1 |
| *fancl* | Fanconi anaemia complementation group L | c63372_g2_i1 | NM_212982.1 |
| Epigenetic regulatory factors | | | |
| *dnmt1* | DNA methyltransferase 1 | c43163_g1_i1 | NM_131189.2 |
| *dnmt3aa* | DNA methyltransferase 3aa | c59097_g4_i2 | NM_001018134.1 |
| Jumonji gene family | | | |

| Gene symbol | Gene description | Contig ID | Reference transcript ID |
|---|---|---|---|
| *jarid2b* | jumonji, AT rich interactive domain 2b | c67175_g1_i2 | NM_001202459.1 |
| *kdm6bb* | lysine (K)-specific demethylase 6B, b | c52506_g1_i1 | NM_001030178.2 |
| Pluripotency factor | | | |
| *pou5f3* | POU domain, class 5, transcription factor 1 | c63041_g1_i1 | NM_131112.1 |

Table S2

(A)

| **RefFinder** | | **Δ CT** | | **BestKeeper** | | | | **NormFinder** | |
|---|---|---|---|---|---|---|---|---|---|
| Genes | GM | Genes | Ave. SD | Genes | SD | Genes | r | Genes | SV |
| *actb1* | 1.19 | *actb1* | 105224.72 | *g6pd* | 600.71 | *g6pd* | 0.42 | *actb1* | 21448.66 |
| *g6pd* | 1.59 | *g6pd* | 125866.30 | *actb1* | 35438.50 | *eef1a1a* | 0.94 | *g6pd* | 93216.54 |
| *eef1a1a* | 3.00 | *eef1a1a* | 188193.69 | *eef1a1a* | 169917.77 | *actb1* | 0.96 | *eef1a1a* | 186876.56 |

(B)

**BestKeeper – Excel-based tool**

| Genes | SD | r | $r^2$ | p-value |
|---|---|---|---|---|
| *g6pd* | 600.71 | 0.42 | 0.18 | 0.003 |
| *actb1* | 35438.50 | 0.96 | 0.92 | 0.001 |
| *eef1a1a* | 169917.77 | 0.94 | 0.88 | 0.001 |
